# A quantitative chemotherapy genetic interaction map reveals new factors associated with PARP inhibitor resistance

**DOI:** 10.1101/171918

**Authors:** Hsien-Ming Hu, Xin Zhao, Swati Kaushik, Lilliane Robillard, Antoine Barthelet, Kevin K. Lin, Andy D. Simmons, Mitch Raponi, Thomas C. Harding, Sourav Bandyopadhyay

## Abstract

Nearly every cancer patient is treated with chemotherapy yet our understanding of factors that dictate response and resistance to such agents remains limited. We report the generation of a quantitative chemical-genetic interaction map in human mammary epithelial cells that charts the impact of knockdown of 625 cancer and DNA repair related genes on sensitivity to 29 drugs, covering all classes of cancer chemotherapeutics. This quantitative map is predictive of interactions maintained in cancer cell lines and can be used to identify new cancer-associated DNA repair factors, predict cancer cell line responses to therapy and prioritize drug combinations. We identify that *GPBP1* loss in breast and ovarian cancer confers resistance to cisplatin and PARP inhibitors through the regulation of genes involved in homologous recombination. This map may help navigate patient genomic data and optimize chemotherapeutic regimens by delineating factors involved in the response to specific types of DNA damage.

## INTRODUCTION

Chemotherapy is given to the vast majority of cancer patients and used based on average responses rather than personalized decisions (Barcenas et al., 2014). Limited improvements in survival by the use of chemotherapy also highlight the need to develop new drugs and make better use of existing drugs (Early Breast Cancer Trialists’ Collaborative, 2005). Furthermore, choosing from multiple possible chemotherapy options can complicate clinical decision making. Therefore, optimizing the use of chemotherapies is a significant and pressing challenge in precision oncology. Chemotherapies commonly target the heightened proliferation resulting from unrestrained cell cycle and DNA damage checkpoints present in cancer cells but their narrow therapeutic window results in the dose-limiting toxicities common with these agents. While tumors that harbor specific alterations in DNA repair genes such as *BRCA1*, *BRCA2* and *ERCC1* are more responsive to certain chemotherapies (Byrski et al., 2012; Olaussen et al., 2006), our knowledge of relevant biomarkers for chemotherapy remains limited. Therefore, understanding the impact that tumor mutations have on modifying drug responses can lead to more efficient use of chemotherapy.

Recent advances in genomics have led to a dramatic increase in the rate of discovery of altered genes in patient tumors. This explosion in knowledge has led to bottlenecks at the level of a functional understanding of tumor genomes, a key step in therapeutic development. Chemical-genetic interaction maps can aid in elucidating roles for genetic events in cancers by causally linking them to drug sensitivity (Martins et al., 2015; Muellner et al., 2011). Furthermore, effectively connecting gene alterations with therapeutics will also require clarity into the exact mechanism of drug action which are often lacking for classical chemotherapeutic agents as well as newly developed drugs targeting DNA repair and processing (Cheung-Ong et al., 2013; Helleday, 2011; Liu et al., 2012; Mitchison, 2012). In the case of PARP inhibitors, their efficacy may be dependent on their ability to trap PARP onto DNA leading to DNA double strand breaks (DSBs) during replication, rather than blocking the repair of single-strand breaks through enzymatic inhibition of PARP as initially hypothesized (Helleday, 2011; Murai et al., 2012). It is likely that insights into the mechanisms of action of chemotherapies will need to be combined with an understanding of gene function in order to create predictive models of drug responses in patients.

A key milestone in the field was the discovery that tumor cells that are deficient in *BRCA1* or *BRCA2* are sensitive to PARP inhibitors in a synthetic lethal manner, ultimately leading to approval of these agents in ovarian cancer. Mechanistically, this synthetic lethal interaction takes advantage of a deficiency in homologous recombination (HR) caused by *BRCA1/2* mutation that is necessary to repair DNA lesions incurred by PARP inhibition (Bryant et al., 2005; Farmer et al., 2005). With the approval of several PARP inhibitors, both *de novo* and acquired resistance to PARP inhibitors has become an important clinical problem. What appears to be critical for resistance is the restoration of HR that in some cases can be attributed to secondary intragenic mutations which restores BRCA1 or BRCA2 functionality (Norquist et al., 2011). Although additional factors have been reported, little is known about their relevance to resistance in the clinic (Lord and Ashworth, 2013). Central to emerging mechanisms of resistance is the interplay between two major repair pathways, non-homologous end joining (NHEJ) and HR. In a competitive model between these two pathways, the NHEJ factor TP53BP1 suppresses HR and TP53BP1 loss restores HR facilitating PARP inhibitor resistance (Bouwman et al., 2010; Bunting et al., 2010; Chapman et al., 2012). However, *TP53BP1* loss has not been observed clinically suggesting additional factors may contribute to the resistant phenotype.

Here, we generate a systematic resource that quantitatively maps the influence of the knockdown of 612 DNA repair and cancer-relevant genes on the responses to 31 chemotherapeutic agents in breast cancer, covering nearly all major FDA approved chemotherapies. We demonstrate that the map recovers many known modulators of chemosensitivity and is able to link therapies with common mechanisms of action. We show that the map is a predictive tool to computationally infer cancer cell line drug sensitivity and design effective drug combinations with inhibitors of the DNA damage signaling kinase ATR. We also identify GPBP1 as a new factor whose loss contributes to PARP inhibitor and platinum resistance, a finding that is supported by data from high grade serous ovarian cancer patients. This chemical-genetic interaction map can be used to identify new factors that dictate responses to chemotherapy and aid in the translation from tumor genomics to therapeutics.

## RESULTS

### Generation of a chemotherapy based genetic interaction map in breast epithelial cells

We developed a quantitative chemical-genetic interaction mapping strategy to uncover the impact of gene loss on proliferative responses to a panel of approved chemotherapies as well as emerging inhibitors of DNA repair. Beyond common tumor suppressor genes, we focused on genes recurrently deleted in breast and ovarian cancer, two diseases which are characterized by a high prevalence focal deletions and amplifications compared to point mutations, providing a rationale for using loss of function genetics (Ciriello et al., 2013). We mined TCGA studies as well as the METABRIC breast cancer cohort covering over 3,000 samples to identify a set of over 200 breast and 170 ovarian cancer genes whose deletion occurred with high frequency in these studies (Figure 1A, Table S1) (Cancer Genome Atlas, 2012; Cancer Genome Atlas Research, 2011; Curtis et al., 2012). We also included nearly all genes known to be involved in DNA repair (n=134). As a complement, we assembled a collection of 29 distinct compounds encompassing nearly all FDA approved chemotherapies for breast and ovarian cancer, four PARP inhibitors and two other targeted therapies (Figure 1B). In addition, we profiled two common drug combinations for a total of 31 distinct treatments. The map was generated in MCF10A cells which are epithelial, derived from healthy breast tissue, devoid of the large numbers of mutations typical of cancer cells, diploid and HR competent (Debnath et al., 2002). By molecular profiling, these cells are receptor-negative or basal-like, a subtype that has been shown to be similar in biology and etiology to high-grade serous ovarian cancer (Cancer Genome Atlas, 2012). Knockdowns were performed using esiRNAs which are enzymatically cleaved long double-stranded RNAs that exist in a pool with high sequence complexity and exhibit less off-target effects and noise than commonly associated with siRNA and shRNA approaches (Kittler et al., 2007). To generate the chemical-genetic interaction map, MCF10A cells were transfected with individual esiRNAs, exposed to either DMSO or drug in parallel, allowed to proliferate for 72 hours before counting. Knockdown of an essential gene, KIF11, was used as positive control for gene knockdown in the screen (Figure S1A). Normalized cell numbers from each knockdown in the presence of drug or DMSO were compared to identify differences in proliferation over 8 replicates (4 in each condition) and the significance of this difference was converted into a signed chemical-genetic interaction score (*S*) (see Methods) (Martins et al., 2015). Positive *S*-scores indicate that the gene loss caused drug resistance and negative *S*-scores indicate that gene loss induced drug sensitivity that could constitute a synthetic lethal interaction. Analysis of the distribution of scores based on knockdown of GFP as negative control allowed the assignment of a false discovery rate (FDR) of 10%, 5% and <1% to cutoffs of *S*=±3, ±4 and ±5, respectively (Figure 1C). Altogether we determined quantitative scores for 19,406 gene-drug interactions and identified 1,042 positive and 740 negative interactions at *S*=±3, corresponding to a 10% FDR (Table S2). These interactions mapped to a median of 27 and 22 positive and negative interactions per drug, respectively (Figure S1B).

**Figure 1:**
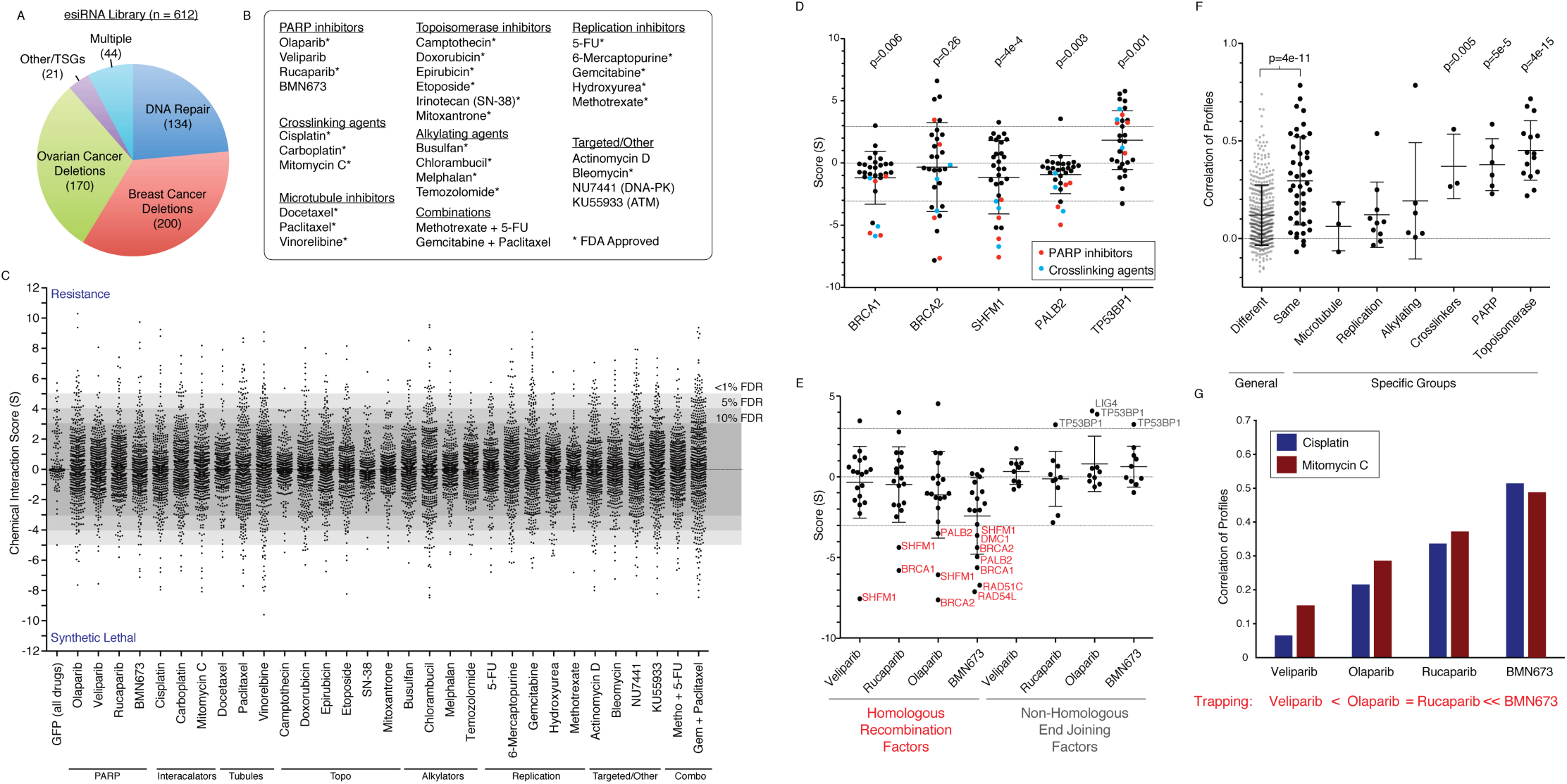
Design of a chemical genetic interaction map and recapitulation of known gene and drug relationships. (A) Composition of genes selected for esiRNA based knockdown. TSG = tumor suppressor genes (B) Selection of 31 drugs profiled in this study. (C) Distribution of chemical genetic interaction scores (S) for all drugs profiled. Scores of 899 GFP knockdowns across all tested drugs are shown. FDR cutoffs are based on the percent of GFP knockdowns falling outside of a given score threshold. Metho = methotrexate, Gem = gemcitabine. (D) Genetic interactions with BRCA-pathway members BRCA1, BRCA2, SHFM1, and PALB2 as well as the NHEJ factor TP53BP1. Interactions with PARP inhibitors and crosslinking agents are highlighted and p-values represent the significance of differences between these scores and zero using a t-test. Dotted lines represent 10% FDR cutoff. (E) PARP inhibitor scores with all 19 annotated HR factors and 10 annotated NHEJ factors. (F) Correlation of chemical-genetic interaction profiles among drugs that are known to belong to the same of different classes. For specific drug classes, pairwise correlations were compared against background of correlations of drugs from different classes and p-value assessed by t-test. (G) Correlation of profiles for PARP inhibitors with two cross linking agents, cisplatin and mitomycin C. Trapping potency from (Murai et al., 2014).

As control, we examined the impact of loss of BRCA proteins on sensitivity to PARP inhibitors, a known synthetic lethal interaction (Bryant et al., 2005; Farmer et al., 2005). Loss of BRCA1 or BRCA2 was among the most synthetic lethal with PARP inhibitors in our dataset including strong interactions with the PARP inhibitor BMN673 (BRCA1 *S*=-4.4, BRCA2 *S*=-5.6). This finding also extended to members of the BRCA pathway, SHFM1 (*S*=-2.9) and PALB2 (*S*=-4.9), which mediate HR as previously reported (Figure 1D) (Buisson et al., 2010; McCabe et al., 2006). We also observed strong synthetic lethal interactions between BRCA1/2 and BRCA-pathway genes and DNA cross-linking agents cisplatin and mitomycin C (BRCA1 with cisplatin, *S*=-5.8, and with mitomycin C, *S*=-5.1) (Figure 1D). In BRCA1 knockout cells, synthetic lethality with PARP inhibitors is related to its ability to trap PARP onto DNA (Murai et al., 2014; Shen et al., 2013). Using the chemical-genetic interaction map we next asked whether this trend extends beyond BRCA1 to the entire HR pathway. We examined known genes involved in HR and found that they were also often synthetic lethal with PARP inhibitors in a manner that was related with the degree of PARP trapping onto DNA (Figure 1E). Illustrating this point, the strongest trapper, BMN673, had an average score of −2.4 with 19 known components of HR (p=3.1e-4), which was less than any other PARP inhibitor. Since these drugs are comparable inhibitors of PARP enzymatic activity, our results indicate that synthetic lethality with loss of components of HR machinery is more dependent on PARP trapping than enzymatic inhibition. Loss of the NHEJ factor TP53BP1 has been shown to cause resistance to PARP inhibitors in several models (Bouwman et al., 2010; Bunting et al., 2010; Chapman et al., 2012). This was also reflected in the chemical-genetic map with TP53BP1 knockdown conferring resistance to PARP inhibitors (BMN673 S=3.3) and DNA cross-linkers (cisplatin S=4.3) (Figure 1D,E). We conclude that the chemical-genetic interaction map recapitulates known drivers of chemo-sensitivity and resistance in a quantitative fashion and is a resource for the identification of potential new drivers of the drug response.

### Chemical-genetic profiles link drugs with similar mechanisms of action

While broad classes of chemotherapeutics target various aspects of DNA processing and repair, their exact mechanisms of action are often unclear (Cheung-Ong et al., 2013). We therefore asked if the map could be used to link together drugs based on common mechanisms of action. For a given drug, its profile of chemical interaction scores represents a high-resolution phenotype that can be compared to other drugs. Calculating all-pairwise correlations between drugs revealed that drugs known to operate in the same general class had a higher average correlation of profiles as compared to drugs that were unrelated (Figure 1F, p=4e-11). Overall, this trend was highest for topoisomerase and PARP inhibitors as well as DNA cross-linkers, which were all significantly more inter-related than compared to background (p<0.05, Figure 1F). For topoisomerase inhibitors, their profiles were highly correlated (mean r=0.45, p=4e-15) and exemplified by the similarity of profiles of topoisomerase II inhibitors, etoposide and doxorubicin (r=0.65, p=5e-79). The ability to link drugs with similar mechanisms of action led us to further investigate the mechanism of action of PARP inhibitors which have been proposed to work through a dual mechanism of enzymatic inhibition as well as trapping of PARP onto DNA during replication (Murai et al., 2012). We found a strong correlation of profiles by comparing PARP inhibitors with cisplatin and mitomycin C that both work by causing intra-strand crosslinks that block replication (average r=0.35, p=6.9e-7). However, this correlation was highly related to PARP trapping ability with the most potent trapper, BMN673, having the highest correlation with cisplatin (r=0.51) and mitomycin C (r=0.49) (Figure 1G). Taken together, our results further support the model whereby PARP trapping creates double strand breaks during replication in a manner similar to cisplatin and mitomycin C and that HR is necessary to repair these lesions. Therefore the genetic interaction map provides a high-resolution means to understand similarities and differences between the mechanism of action of drugs.

### Prediction of cancer cell line responses using the chemical-genetic interaction map

Based on the similarity of profiles between related drugs we next sought to combine genetic interactions based on drug class to identify a consensus chemical-genetic interaction map. In this consensus map a connection between a gene and a compound category reflects a concordance of response across multiple related drugs and compared against a randomly permuted background. At an FDR of 0.1% we identified 125 connections between genes and different drug classes (Figure 2A, Table S3). While connections spanned all major drug classes, topoisomerase inhibitors, PARP inhibitors and alkylating agents made up the majority of this network while microtubule inhibitors were under-represented due to the lack of genetic interactions in common across this class of agents (Figure S1C,D). Through the analysis of independent chemical entities sharing a common mechanism this map highlights many potentially new modifiers of drug responses that are altered in breast and ovarian cancers that may participate in the DNA damage response.

**Figure 2:**
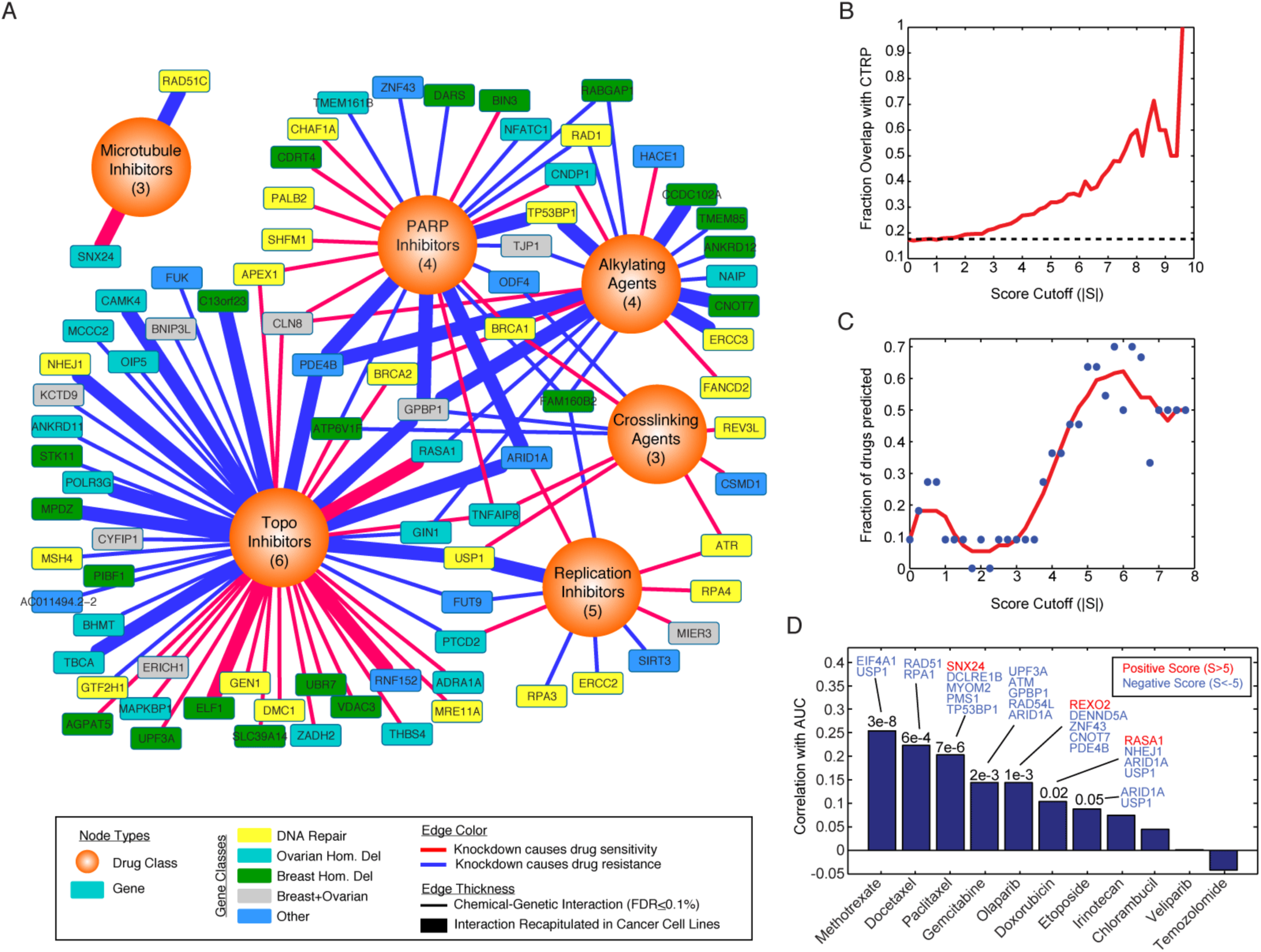
Prediction of cell lines responses from the chemical-interaction map. (A) Consensus interaction map based on coordinate responses with drug classes. All interactions shown have a FDR of category association <0.1%. The number of drugs in each category is indicated. Thicker edges represent interactions that are also found across cancer cell line collections (p<0.01). (B) Overlap with correlation based chemical-genetic interactions from cancer cell lines. Shown is the fraction of chemical genetic interactions at a given score cutoff (|S|) where the expression of the gene is also significantly associated with resistance or sensitivity to the same drug across cell lines in the CTRP dataset (p<0.01). Dotted line represents baseline overlap at random (17.3%). (C) Prediction of cell line responses to 11 drugs overlapping with the CTRP dataset. Cell lines were scored based on the sum of normalized gene expression for all genes in the network at a given cutoff where the contribution of each gene to the sum is weighted positively or negatively based on the sign in the chemical-genetic interaction map. These drug and cell line specific scores are then correlated with the AUC values reported in the CTRP and significant predictors are counted (p≤0.05). Red line is a sliding average. (D) Analysis of cell line response predictions based on a score cutoff of 5. For each model the correlation of predicted versus real AUC is shown with accompanying p-value when significant. Genes whose expression contributed the most to the prediction accuracy are shown (see Methods).

The ability of the chemical-genetic interaction map to identify causal genetic relationships also raises the possibility that a quantitative map could complement pharmacogenomics efforts using large panels of cancer cell lines (Barretina et al., 2012; Basu et al., 2013; Garnett et al., 2012). While previous studies have used supervised machine learning approaches to identify molecular correlates of drug sensitivity across cell lines, we hypothesized that the relationships identified by gene knockdown constitute a more direct and causal readout of gene function that could enhance biomarker identification. Comparison of 11 drugs in common between our study and the Cancer Therapeutics Response Portal (CTRP) revealed a strong degree of overlap between interactions identified in the chemical genetic interaction map and genes whose response was significantly correlated with the drug response (Figure 2B) (Basu et al., 2013). Furthermore, this degree of overlap was highly related to the score threshold used with 21.5% of interactions overlapping at a cutoff of 3 (p=2.9e-3) and nearly 60% overlapping at a cutoff of 8 (p=3.1e-5), reflecting the quantitative nature of the dataset (Figure 2B).

The significance and quantitative nature of the overlap between our map and expression based correlates of drug sensitivity found in cancer cell lines prompted us to explore whether this map could be used to systematically predict cancer cell line sensitivity in an unsupervised fashion. For each drug we used the relative expression of each of the genes in its network to derive a drug response prediction for every cell line (see Methods). We evaluated this method using a sliding cutoff to define the specific network for each drug and found that more stringent networks provided increased power to predict drug sensitivity with nearly 60-70% of drugs predicted accurately at a cutoff between 5 and 6 (Figure 2C). At a cutoff of 5, predictions were significant for 7 out of the 11 drugs (Figure 2D). Analysis of the genes that were most informative in making correct predictions in these cases revealed novel genes involved in drug sensitivity and resistance. Knockdown of EIF4A1 caused resistance to methotrexate (S=6.5) and in cell lines EIF4A1 expression is positively correlated with methotrexate sensitivity across 645 cell lines (r=0.25, p=1.9e-8), consistent with the network prediction. Alternatively, SNX24 knockdown was synthetic lethal with paclitaxel (S=-6.8) and SNX24 expression was negatively correlated with drug sensitivity (r=-0.14, p=0.0026). Thus computational analysis of chemical-genetic interaction maps can be used to complement cancer cell line screens and may be able to produce biomarkers that bridge correlation with causation.

### Prediction of drug synergies using the chemical-genetic interaction map

There has been considerable interest in the development of targeted therapies that inhibit DNA repair machinery to be used in combination with agents that generate specific types of DNA damage (Gavande et al., 2016). We observed that loss of the DNA damage signaling kinase ATR induced sensitivity to crosslinking agents and inhibitors of DNA replication in the consensus map (Figure 2A). This was in contrast to its closely related paralog kinase ATM, which was not linked to the response to these drugs but rather with moderate resistance to DNA alkylators and nucleotide analogs (Table S3). These results reflect the importance of ATR in the repair of damage during DNA replication as opposed to ATM (Flynn and Zou, 2011). We hypothesized that the synthetic lethal interactions observed with ATR knockdown could be phenocopied with a small molecule inhibitor of ATR and used to prioritize synergistic drug combinations. We tested the combined effects of the ATR inhibitor VE-821 with five drugs that were synthetic lethal with ATR knockdown and three drugs that were not (Figure 3A). Using a matrix screening approach we measured the effects of each drug on proliferation and determined a synergy score reflecting the degree of drug synergy based on the Loewe excess model (Lehar et al., 2009). Drugs that were synthetic lethal with ATR inhibition were more synergistic than those that were not predicted to be (p=0.048, Figure 3B). In particular, for cisplatin and BMN673 we observed marked synergy at multiple doses, in short as well as long-term growth assays (Figure 3C-E). Since little is known about ATR and PARP inhibitor synergy in breast cancer, we explored the degree to which this interaction could be recapitulated in breast cancer cell lines. We found that the addition of VE-821 was synergistic and could potentiate the activity of BMN673 in 3 out of the 5 cell lines we tested (60%) with a combination index (CI) between 0.07-0.17 with CI < 0.3 indicating strong synergy (Figure 3F) (Chou, 2006). Therefore the chemical genetic interaction map can be used to prioritize new drug combinations and in the case of ATR inhibitors, highlights drugs combinations that could help optimize clinical testing.

**Figure 3:**
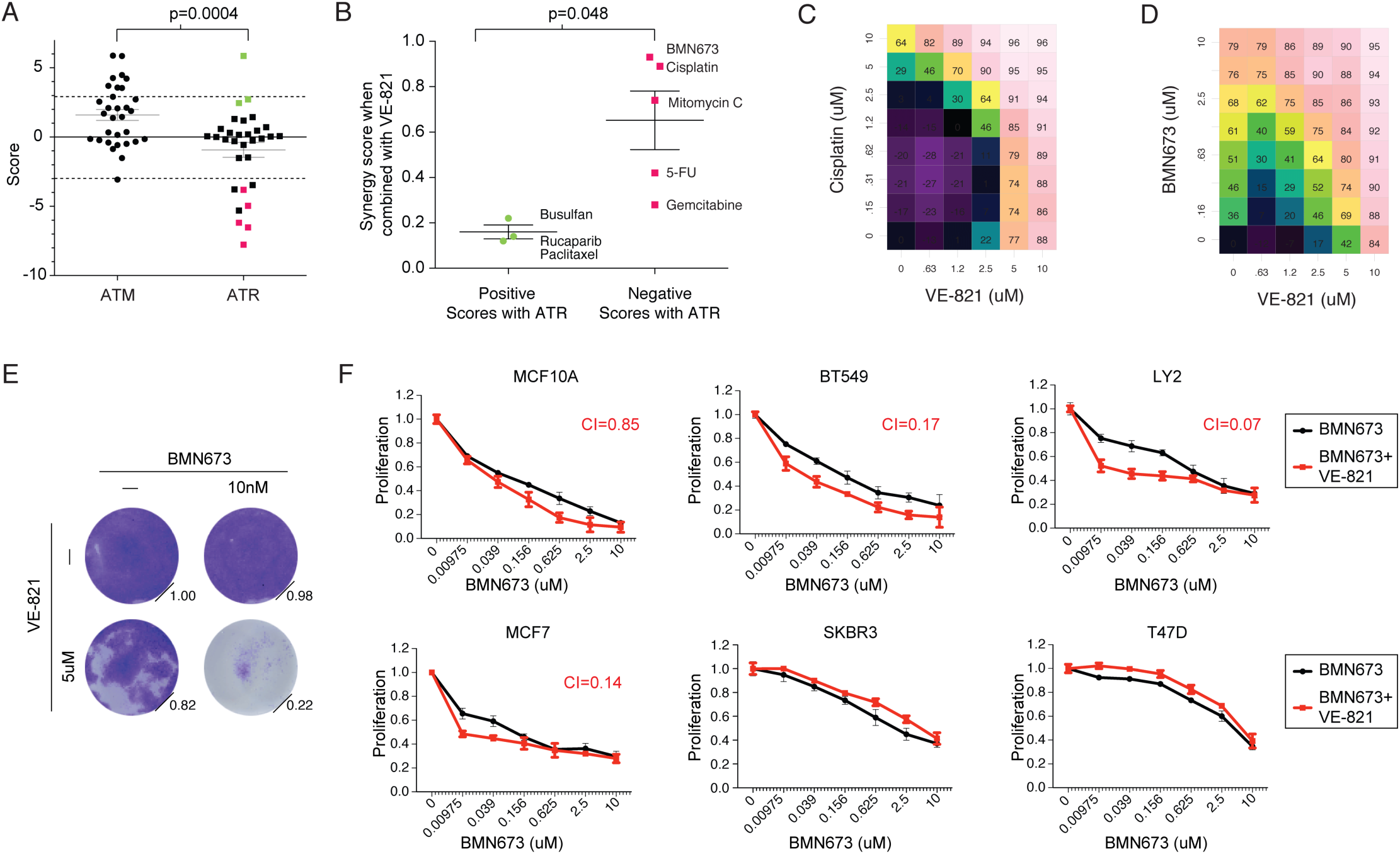
Prediction of drug synergies based on ATR inhibition. (A) Comparison of chemical interaction scores for ATM and ATR knockdown. P-value based on difference in overall score distribution. Selected positive (green) and negative (red) drugs for ATR are shown. (B) Synergy scores between the ATR inhibitor VE-821 and drugs selected based on positive and negative scores with ATR in the map. Percent inhibition of growth of MCF10A cells treated with an escalating dose matrix of VE-821 with (C) cisplatin and (D) BMN673. (E) Crystal violet staining of MCF10A cells 14 days after treatment with vehicle, VE-821, BMN673 and the combination. Normalized quantification of intensity shown. (F) Relative proliferation of MCF10A and indicated breast cancer cell lines treated with BMN673 alone versus both BMN673 and VE821 together for 3 days. Single agent BMN673 was normalized to DMSO and combinations were normalized to VE-821 alone. A constant dose of 2.5uM VE-821 was used except in MCF7 cells where 625nM was used due to the toxicity of single agent VE-821 at higher doses. The combination index (CI) value at the indicated doses is shown. (A,B) Error bars are s.e.m.

### Prediction of factors mediating resistance to PARP inhibition

We next evaluated the map as a systematic resource to predict new molecules involved in DNA repair and new mechanisms of resistance to chemotherapy. We focused on PARP inhibitors olaparib, veliparib, rucaparib and BMN673 as well as cisplatin since they are of high clinical interest, have similar mechanisms of action and generate DNA damage that depends on repair via HR (Figure 1E). As controls, BRCA1, BRCA2, PALB2 and SHFM1 knockdown was synthetic lethal with these agents and loss of TP53BP1 was associated with resistance (Figure 4A). An important consideration in interpreting genetic interaction data from a single cell line is the degree to which such interactions are maintained in other cellular contexts (Ashworth et al., 2011). To assess this we identified chemical-genetic interactions using the same experimental approach with BMN673 in two breast cancer lines, MDAMB231 and SUM149PT, and two ovarian cancer lines OVCAR3 and UWB1.289. After scoring esiRNAs for their ability to induce resistance or sensitivity to BMN673 in each of these lines (defined based on a cutoff of p<0.01), we found that the chemical-genetic interaction score was highly predictive of whether a particular interaction was preserved in other cell lines (Table S4). For example, interactions with BMN673 that had a score >5 in MCF10A cells were 40% likely to validate in 2 or more cell lines and 70% likely to validate in at least one other line (Figure 4B). Although this trend was also evident for negative interactions, the magnitude was much less pronounced. Since two of the tested lines were *BRCA1* mutant (SUM149PT, UWB1.289), a likely reason for this difference is that factors whose loss leads to PARP sensitivity in HR-competent MCF10A cells may not be relevant in *BRCA1* mutant cells that are already HR deficient. Therefore, interaction strength in MCF10A cells can be used to predict genetic interactions in other cell lines highlighting the quantitative nature of this map.

**Figure 4:**
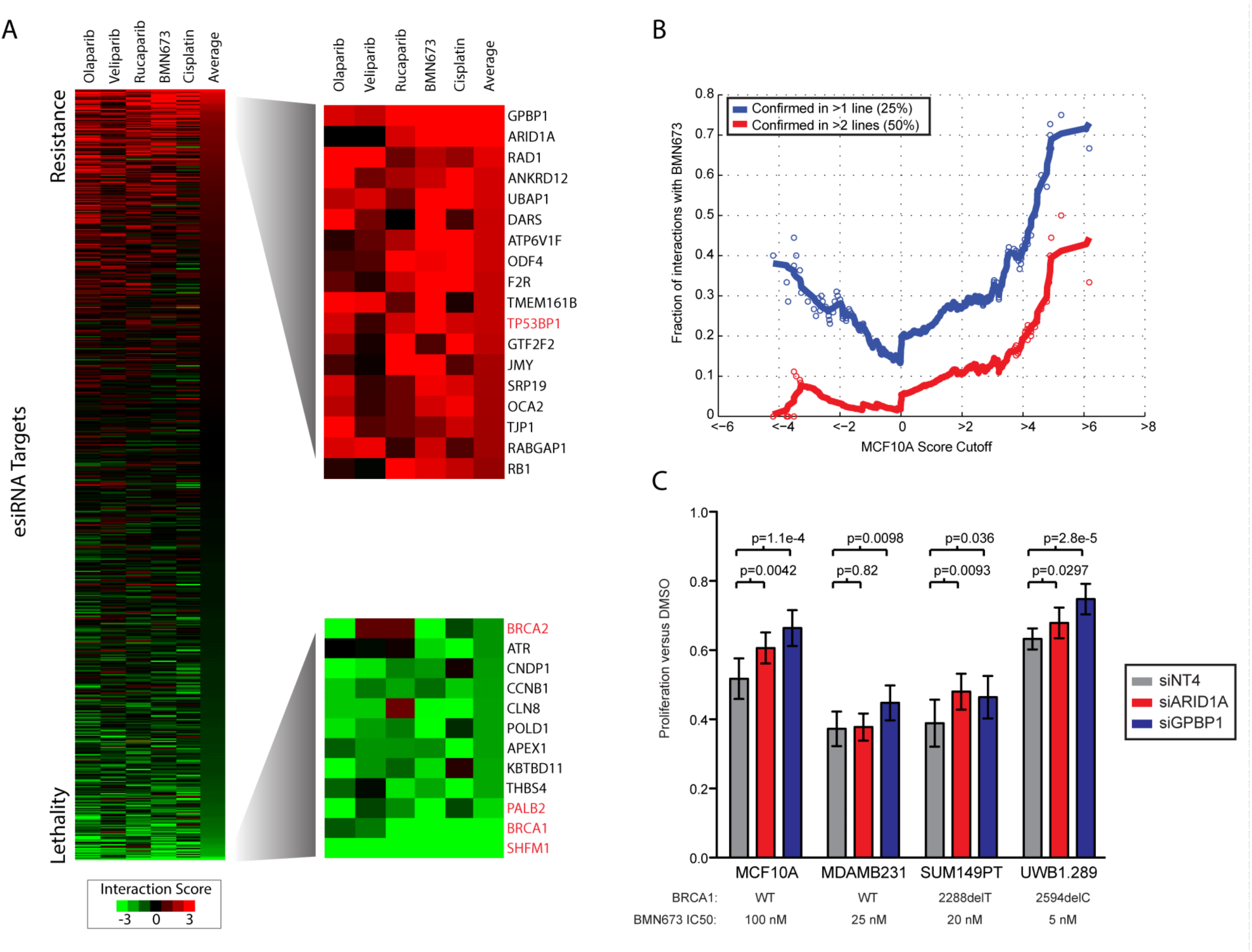
Assessment of genetic interactions with PARP inhibitors and cisplatin. (A) Interaction profiles of four PARP inhibitors and cisplatin sorted based on average across all drugs. Known factors associated with resistance and sensitivity indicated in red. (B) Preservation of genetic interactions with BMN673 between MCF10A cells and four independent cancer cell lines, MDAMB231, SUM149PT, OVCAR-3 and UWB1.289. A genetic interaction is considered preserved if it is significant (p<0.01) with the same direction in one or more lines. Each point represents the cumulative rate of preservation in >1 line (blue) or >2 lines (red) for all interactions scoring past a particular cutoff. Solid lines are sliding averages. (C) Confirmation of resistance interactions using independent synthetic siRNA gene knockdown in cell lines. Samples were treated with approximate IC_50_ dose of BMN673 and normalized to gene knockdown treated with DMSO. NT4 is non-targeting control.

We next sought to validate the top two hits producing resistance, GPBP1 and ARID1A in additional models. Using independent siRNAs we confirmed that loss of either factor caused resistance to BMN673 in MCF10A, MDAMB231, SUM149PT and UWB1.289 cells in most, if not all, cases (Figure 4C). ARID1A loss often occurs through somatic mutation and has been previously linked to the regulation of DNA repair processes (Dykhuizen et al., 2013; Shen et al., 2015). We next confirmed this result in engineered ARID1A -/- MCF10A cells, which were resistant to BMN673 in comparison to parental MCF10A (Figure S2A). We next searched for clinical evidence that ARID1A loss contributes to resistance to PARP or platinum containing chemotherapy. In support, we found ARID1A loss was linked to poor outcome in TCGA high-grade serous ovarian cancers (TCGA HGSOC) receiving platinum as standard of care (p=0.01, Figure S2B) (Cancer Genome Atlas Research, 2011). To test if this observation extended to PARP inhibitors, we analyzed samples from high-grade serous or endometrioid ovarian cancer patients treated with rucaparib (NCT01891344) (Swisher et al., 2017). We did not identify any patients with concurrent *BRCA1* and *ARID1A* mutations and therefore focused our analysis on a cohort of 154 patients that did not have known mutations in the HR pathway where we identified 10 that had mutations in ARID1A. The progression free survival (PFS) for these 10 *ARID1A* mutant cases was significantly lower than for the rest of this cohort (p=0.003, Figure S2C). These clinical data show that PARP inhibitors provide no clinical benefit in ARID1A mutated high-grade serous or endometrioid ovarian cancer and warrants further investigation.

### GPBP1 loss causes PARP and platinum resistance by regulating the expression of factors involved in homologous recombination

We next investigated the top candidate in our categorical analysis, GPBP1, a transcription factor of unknown function. GPBP1 lies on chromosome 5q11, a region focally deleted in approximately 5% of TCGA HGSOC and 4% of TCGA breast cancers. To determine if GPBP1 plays a role in the transcriptional response to DNA damage, we performed RNAseq analysis of control and GPBP1-knockdown MCF10A cells treated with or without BMN673. RT-qPCR of GPBP1 knockdown cells confirmed 90% knockdown at the mRNA level in this experiment (Figure S3A). In response to 24 hour BMN673 treatment, GPBP1 knockdown caused the up-regulation of factors involved in HR based on gene set enrichment analysis (GSEA) (Subramanian et al., 2005) (Figure 5A), indicating a potential compensatory mechanism to facilitate repair of lesions incurred by PARP inhibition. In contrast to control cells, GPBP1 knockdown resulted in up-regulation of distinct and canonical HR factors such as BRCA1 and RAD51B in response to BMN673 (Figure 5B).

**Figure 5:**
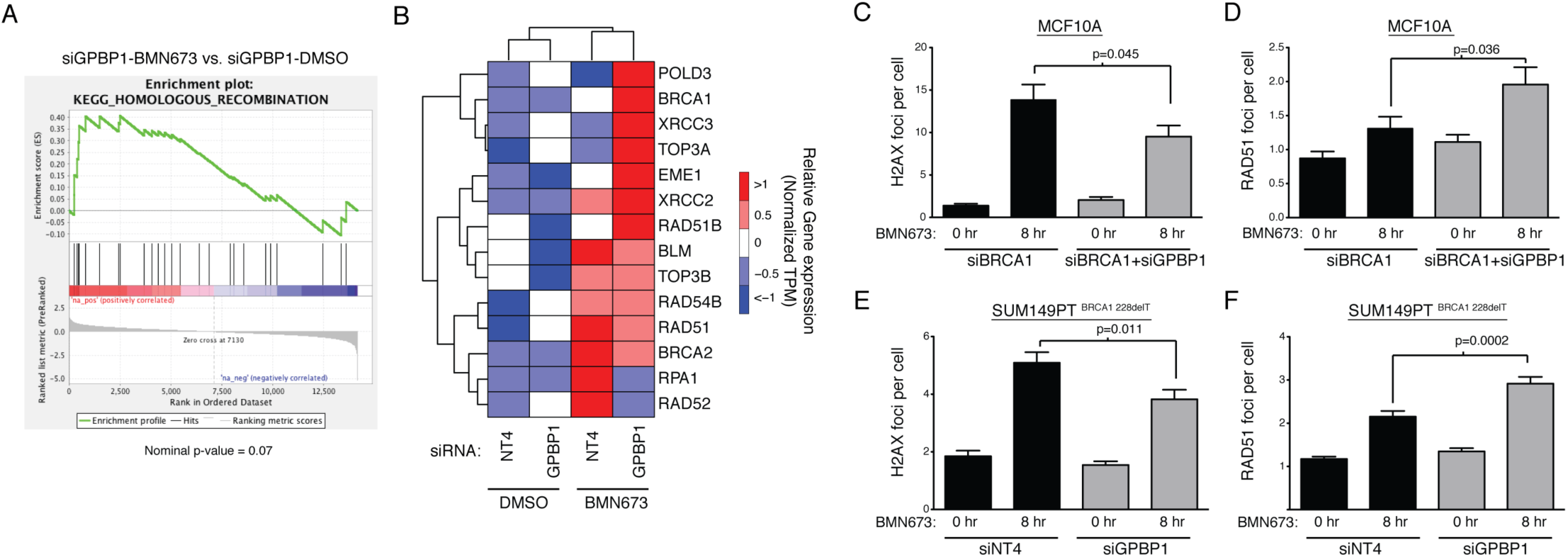
BMN673 treatment of GPBP1 knockdown cells causes up regulation of the homologous recombination pathway. (A) Gene set enrichment analysis of homologous recombination pathway genes using RNAseq data from MCF10A cells with GPBP1 knockdown in presence and absence of BMN673. (B) Heatmap representation of expression of HR pathway genes differentially expressed in the presence of BMN673. TPM = transcripts per kilobase million. (C) Quantification of gamma-H2AX foci and (D) RAD51 recruitment after treatment with 500nM of BMN673 in the presence of the indicated gene knockdowns in MCF10A cells. (E) Quantification of gamma-H2AX foci and (F) RAD51 recruitment after treatment with 50nM of BMN673 in the presence of the indicated gene knockdowns in SUM149PT cells. NT4 is non-targeting control. Error bars s.e.m.

We next asked if this transcriptional response was sufficient to enhance the repair of double strand breaks incurred by PARP inhibition and if this occurred via HR. This hypothesis was particularly intriguing since GPBP1 knockdown caused resistance to BMN673 in *BRCA1* mutant cancer cell lines suggesting that GPBP1 loss may bypass the requirement of BRCA1 for HR (Figure 4C). To test this hypothesis we established a HR-deficient and PARP inhibitor sensitive MCF10A model by BRCA1 knockdown and in this model knockdown of BRCA1 and GPBP1 together led to a significant rescue of BMN673 sensitivity (Figure S3B). Using immunofluorescence we found that generation of γH2AX foci after BMN673 treatment was reduced in BRCA1+GPBP1 versus BRCA1 knockdown cells (p=0.045), indicating that GPBP1 loss led to a reduction in the number of DNA double strand breaks formed after PARP inhibitor treatment (Figure 5C). To determine if this reduction in double strand breaks was due to heightened HR repair capacity we examined the recruitment of the strand-exchange protein RAD51 to damaged chromatin, a mark of commitment to double strand break repair using HR. We found that the recruitment of RAD51 was increased in BRCA1+GPBP1 versus BRCA1 knockdown cells after PARP inhibition indicating that GPBP1 loss led to an increase in double strand break repair through HR (p=0.036, Figure 5D). We confirmed these findings in *BRCA1* mutant SUM149PT cells where GPBP1 knockdown also led to a significant reduction in H2AX foci and increase in RAD51 foci after BMN673 treatment (Figure 5E,F) indicating that GPBP1 loss can also restore HR in cases when *BRCA1* is mutated. These results indicate that GPBP1 loss contributes to increased double strand break repair by HR as a mechanism of PARP inhibitor resistance.

We next asked if the expression of HR factors was also elevated in human cancer samples harboring GPBP1 loss and if it might contribute to drug resistance. There was a strong concordance between genes up-regulated upon GPBP1 knockdown and genes whose expression level was higher in breast cancers with GPBP1 loss (Figure 6A). Further analysis of samples with GPBP1 loss in TCGA ovarian cancer samples also reflected the increased expression of a number of the same HR factors, indicating a similar function of GPBP1 in these two disease types (Figure 6B). We next asked if this enhancement in HR gene expression upon GPBP1 loss resulted in drug resistance in ovarian cancer patients treated with platinum containing therapy. In the TCGA ovarian cohort, survival analysis indicated that GPBP1 loss was associated with poor outcome and resistance to platinum therapy (p=0.001 via log-rank test, Figure 6C). Therefore GPBP1 loss contributes to platinum resistance in ovarian cancer through the increased expression of genes involved in HR.

**Figure 6:**
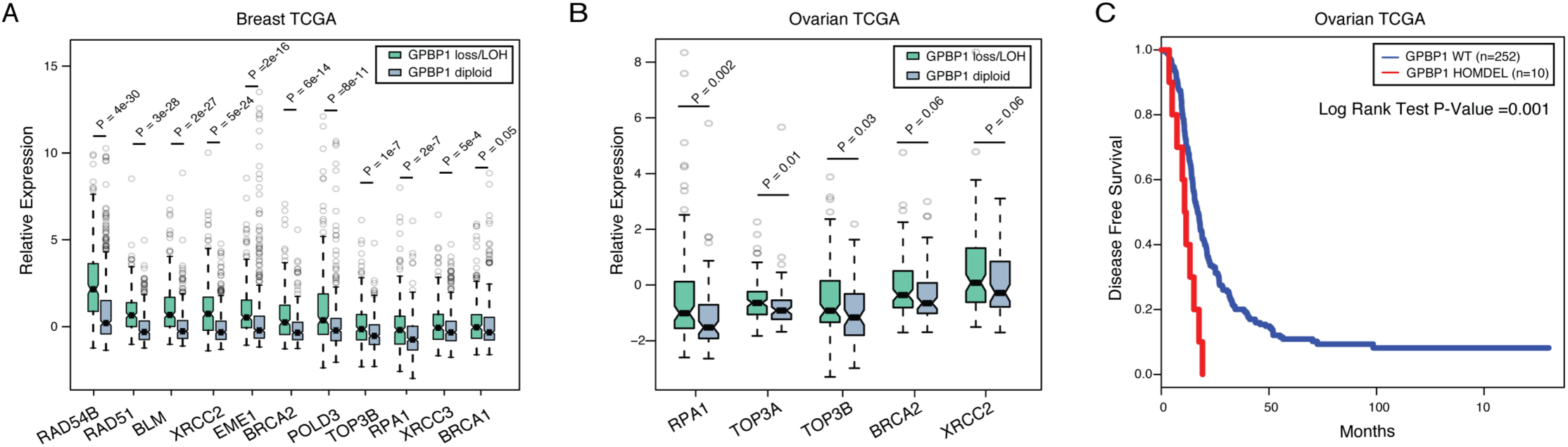
Cancers with GPBP1 loss display heightened expression of HR genes and resist platinum treatment in ovarian cancer. (A) Comparison of gene expression levels of homologous recombination pathway genes in the breast TCGA among tumors with GPBP1 homozygous/heterozygous loss versus diploid CNV status. (B) Comparison of gene expression levels of homologous recombination pathway genes in ovarian cancer patients from TCGA High Grade Serous Ovarian Cancer (HGSOC) dataset with GPBP1 homozygous/heterozygous loss versus diploid CNV status. P-values were calculated by non-parametric Mann-Whitney-Wilcoxon test. (C) The Kaplan-Meier disease free survival (DFS) analysis of patients in TCGA HGSOC with samples with deletion in GPBP1. Boxes represent the interquartile range and whiskers are 1.5 times the interquartile range.

## DISCUSSION

We present a quantitative map to link the efficacy of chemotherapeutics to tumor genetics. This dataset and general approach can serve as a platform for the functional and therapeutic translation of tumor genomes. In contrast to most standard genetic screens, this approach provides a quantitative readout which approximates genetic interaction strength and allows for the comparison of responses across many drugs. Covering nearly 20,000 interactions, this map is larger than previously published mammalian chemical-genetic interaction maps by nearly an order of magnitude (Martins et al., 2015). To aid in integration of these data with ongoing efforts to systematize cancer related screens, data from this network have been deposited into the Cancer Target Discovery and Development (CTD^2^) dashboard (https://ctd2-dashboard.nci.nih.gov/).

Using insights derived from the chemical-genetic interaction map, we highlight several vignettes describing how it can be used to aid in the development of cancer therapeutics. The map was able to identify drugs with similar mechanisms of action and highlights the commonalities between PARP inhibitors and DNA cross-linking agents that contribute to synthetic lethality with loss of HR pathway genes. Computational analysis of the map was used to predict the sensitivity of tumor cells to chemotherapies as well as predict synergistic drug combinations. For example, by tracing drug sensitivities associated with ATR knockdown, we predicted and tested synergy between the ATR inhibitor VE-821 and PARP inhibitor BMN673 in breast cancer cells. Systematic testing indicated synergy in most breast cancer cell lines and future work may evaluate what dictates synergy in breast cancer and other disease subsets that have been reported (Kim et al., 2017; Mohni et al., 2015). In addition, the map may provide a platform for enhancing methods to predict drug responses from baseline genomic profiles, since current approaches do not already use this functional information (Costello et al., 2014). Therefore, this work provides the conceptual framework to incorporate chemical-genetic interaction maps into more direct drug development approaches.

We demonstrate several ways to enhance the reliability and utility of this map. First, we show that related drugs have similar genetic interaction profiles and that analyzing multiple compounds with the same mechanism of action together the map can be used to identify new modifiers of therapeutic responses that are not specific to a single compound. As specific drugs may have unique off targets, such as the case for PARP inhibitors (Knezevic et al., 2016), analyzing related drugs together may identify genetic interactions linked to their core mechanism of action. Second, the plasticity in genetic networks has been an impediment to the identification of genetic interactions that are cell type independent (i.e. ‘hard’ versus ‘soft’ interactions) (Ashworth et al., 2011). Rescreening in multiple cancer cell lines showed that the strength of genetic interaction was proportional to the likelihood of interaction being conserved in other cell lines. Therefore our data indicate that the quantitative nature of genetic interaction maps could be used to distinguish between interactions that are more globally preserved versus those more specific to the cell line tested.

Based on our categorical analysis, we identified that ARID1A loss causes PARP inhibitor resistance in our models. Concordant with our pre-clinical and clinical data, low ARID1A expression has been linked with poor outcome and platinum resistance in high grade serous ovarian cancer (Yokoyama et al., 2014) and clear cell ovarian cancers (Itamochi et al., 2015; Katagiri et al., 2012). However, the functional role of ARID1A on DNA repair is unclear with conflicting reports of its role in homologous recombination (Adamson et al., 2012; Shen et al., 2015). Together, these data warrant a more complete interrogation of the role of ARID1A on PARP inhibitor resistance. The strongest resistance interaction with PARP inhibitors and cisplatin was GPBP1 that is involved in the transcriptional regulation of genes involved in HR. Another transcriptional regulator, CDK12 has been shown to modulate PARP inhibitor sensitivity through the regulation of genes involved in HR (Bajrami et al., 2014; Johnson et al., 2016). Future studies may seek to identify the potential interplay between the targets of CDK12 and GPBP1. Since we observed that GPBP1 loss is linked to chemoresistance and poor clinical outcome, these functional and clinical data warrant a more complete interrogation of the role of GPBP1 and its role in chemoresistance. For example, although GPBP1 loss was not assessed in our rucaparib clinical trial cohort, future work could determine its clinical association with PARP inhibitor resistance. This work highlights the utility of a systematic chemical-genetic interaction map as a resource for the identification of clinically relevant biomarkers of drug susceptibility as well as a foundation for integration with other cancer datasets to enhance drug and biomarker development.

## AUTHOR CONTRIBUTIONS

H.H., S.B. and T.H. conceived the study. H.H., X.Z., S.K., L.R., A.B., K.L., S.B. generated data and participated in computational analyses. A.S, M.R., T.H. and S.B. administered the project. S.B. supervised the study. All authors wrote and approved the manuscript.

## ACKNOWLEDGMENTS

This work was funded by NCI U01CA168370, the UCSF Program in Breakthrough Biomedical Research (PBBR) and the UCSF Beast Oncology SPORE development award to S.B. We thank members of the Bandyopadhyay Lab and Sophia Pfister for helpful comments and critical review of the manuscript.

## Supplementary Tables

Table S1: List of genes with annotations used in the primary screen and drugs and concentrations used.

Table S2: Chemical genetic interaction scores.

Table S3: List of scores based on drug class analysis

Table S4: Individual cell line results from the BMN673 rescreen.

## REFERENCES

Adamson, B., Smogorzewska, A., Sigoillot, F.D., King, R.W., and Elledge, S.J. (2012). A genome-wide homologous recombination screen identifies the RNA-binding protein RBMX as a component of the DNA-damage response. Nature cell biology 14, 318-328.

Ashworth, A., Lord, C.J., and Reis-Filho, J.S. (2011). Genetic interactions in cancer progression and treatment. Cell 145, 30-38.

Bajrami, I., Frankum, J.R., Konde, A., Miller, R.E., Rehman, F.L., Brough, R., Campbell, J., Sims, D., Rafiq, R., Hooper, S., et al. (2014). Genome-wide profiling of genetic synthetic lethality identifies CDK12 as a novel determinant of PARP1/2 inhibitor sensitivity. Cancer research 74, 287-297.

Barcenas, C.H., Niu, J., Zhang, N., Zhang, Y., Buchholz, T.A., Elting, L.S., Hortobagyi, G.N., Smith, B.D., and Giordano, S.H. (2014). Risk of hospitalization according to chemotherapy regimen in early-stage breast cancer. Journal of clinical oncology : official journal of the American Society of Clinical Oncology 32, 2010-2017.

Barretina, J., Caponigro, G., Stransky, N., Venkatesan, K., Margolin, A.A., Kim, S., Wilson, C.J., Lehar, J., Kryukov, G.V., Sonkin, D., et al. (2012). The Cancer Cell Line Encyclopedia enables predictive modelling of anticancer drug sensitivity. Nature 483, 603-607.

Basu, A., Bodycombe, N.E., Cheah, J.H., Price, E.V., Liu, K., Schaefer, G.I., Ebright, R.Y., Stewart, M.L., Ito, D., Wang, S., et al. (2013). An interactive resource to identify cancer genetic and lineage dependencies targeted by small molecules. Cell 154, 1151-1161.

Bouwman, P., Aly, A., Escandell, J.M., Pieterse, M., Bartkova, J., van der Gulden, H., Hiddingh, S., Thanasoula, M., Kulkarni, A., Yang, Q., et al. (2010). 53BP1 loss rescues BRCA1 deficiency and is associated with triple-negative and BRCA-mutated breast cancers. Nature structural & molecular biology 17, 688-695.

Bryant, H.E., Schultz, N., Thomas, H.D., Parker, K.M., Flower, D., Lopez, E., Kyle, S., Meuth, M., Curtin, N.J., and Helleday, T. (2005). Specific killing of BRCA2-deficient tumours with inhibitors of poly(ADP-ribose) polymerase. Nature 434, 913-917.

Buisson, R., Dion-Cote, A.M., Coulombe, Y., Launay, H., Cai, H., Stasiak, A.Z., Stasiak, A., Xia, B., and Masson, J.Y. (2010). Cooperation of breast cancer proteins PALB2 and piccolo BRCA2 in stimulating homologous recombination. Nature structural & molecular biology 17, 1247-1254.

Bunting, S.F., Callen, E., Wong, N., Chen, H.T., Polato, F., Gunn, A., Bothmer, A., Feldhahn, N., Fernandez-Capetillo, O., Cao, L., et al. (2010). 53BP1 inhibits homologous recombination in Brca1-deficient cells by blocking resection of DNA breaks. Cell 141, 243-254.

Byrski, T., Dent, R., Blecharz, P., Foszczynska-Kloda, M., Gronwald, J., Huzarski, T., Cybulski, C., Marczyk, E., Chrzan, R., Eisen, A., et al. (2012). Results of a phase II open-label, non-randomized trial of cisplatin chemotherapy in patients with BRCA1-positive metastatic breast cancer. Breast Cancer Res 14, R110.

Cancer Genome Atlas, N. (2012). Comprehensive molecular portraits of human breast tumours. Nature 490, 61-70.

Cancer Genome Atlas Research, N. (2011). Integrated genomic analyses of ovarian carcinoma. Nature 474, 609-615.

Chapman, J.R., Sossick, A.J., Boulton, S.J., and Jackson, S.P. (2012). BRCA1-associated exclusion of 53BP1 from DNA damage sites underlies temporal control of DNA repair. Journal of cell science 125, 3529-3534.

Cheung-Ong, K., Giaever, G., and Nislow, C. (2013). DNA-damaging agents in cancer chemotherapy: serendipity and chemical biology. Chemistry & biology 20, 648-659.

Chou, T.C. (2006). Theoretical basis, experimental design, and computerized simulation of synergism and antagonism in drug combination studies. Pharmacological reviews 58, 621-681.

Ciriello, G., Miller, M.L., Aksoy, B.A., Senbabaoglu, Y., Schultz, N., and Sander, C. (2013). Emerging landscape of oncogenic signatures across human cancers. Nature genetics 45, 1127-1133.

Costello, J.C., Heiser, L.M., Georgii, E., Gonen, M., Menden, M.P., Wang, N.J., Bansal, M., Ammad-ud-din, M., Hintsanen, P., Khan, S.A., et al. (2014). A community effort to assess and improve drug sensitivity prediction algorithms. Nature biotechnology 32, 1202-1212.

Curtis, C., Shah, S.P., Chin, S.F., Turashvili, G., Rueda, O.M., Dunning, M.J., Speed, D., Lynch, A.G., Samarajiwa, S., Yuan, Y., et al. (2012). The genomic and transcriptomic architecture of 2,000 breast tumours reveals novel subgroups. Nature 486, 346-352.

Debnath, J., Mills, K.R., Collins, N.L., Reginato, M.J., Muthuswamy, S.K., and Brugge, J.S. (2002). The role of apoptosis in creating and maintaining luminal space within normal and oncogene-expressing mammary acini. Cell 111, 29-40.

Dykhuizen, E.C., Hargreaves, D.C., Miller, E.L., Cui, K., Korshunov, A., Kool, M., Pfister, S., Cho, Y.J., Zhao, K., and Crabtree, G.R. (2013). BAF complexes facilitate decatenation of DNA by topoisomerase IIalpha. Nature 497, 624-627.

Early Breast Cancer Trialists’ Collaborative, G. (2005). Effects of chemotherapy and hormonal therapy for early breast cancer on recurrence and 15-year survival: an overview of the randomised trials. Lancet 365, 1687-1717.

Farmer, H., McCabe, N., Lord, C.J., Tutt, A.N., Johnson, D.A., Richardson, T.B., Santarosa, M., Dillon, K.J., Hickson, I., Knights, C., et al. (2005). Targeting the DNA repair defect in BRCA mutant cells as a therapeutic strategy. Nature 434, 917-921.

Flynn, R.L., and Zou, L. (2011). ATR: a master conductor of cellular responses to DNA replication stress. Trends in biochemical sciences 36, 133-140.

Garnett, M.J., Edelman, E.J., Heidorn, S.J., Greenman, C.D., Dastur, A., Lau, K.W., Greninger, P., Thompson, I.R., Luo, X., Soares, J., et al. (2012). Systematic identification of genomic markers of drug sensitivity in cancer cells. Nature 483, 570-575.

Gavande, N.S., VanderVere-Carozza, P.S., Hinshaw, H.D., Jalal, S.I., Sears, C.R., Pawelczak, K.S., and Turchi, J.J. (2016). DNA repair targeted therapy: The past or future of cancer treatment? Pharmacology & therapeutics 160, 65-83.

Helleday, T. (2011). The underlying mechanism for the PARP and BRCA synthetic lethality: clearing up the misunderstandings. Molecular oncology 5, 387-393.

Itamochi, H., Oumi, N., Oishi, T., Shoji, T., Fujiwara, H., Sugiyama, T., Suzuki, M., Kigawa, J., and Harada, T. (2015). Loss of ARID1A expression is associated with poor prognosis in patients with stage I/II clear cell carcinoma of the ovary. International journal of clinical oncology 20, 967-973.

Johnson, S.F., Cruz, C., Greifenberg, A.K., Dust, S., Stover, D.G., Chi, D., Primack, B., Cao, S., Bernhardy, A.J., Coulson, R., et al. (2016). CDK12 Inhibition Reverses De Novo and Acquired PARP Inhibitor Resistance in BRCA Wild-Type and Mutated Models of Triple-Negative Breast Cancer. Cell reports 17, 2367-2381.

Katagiri, A., Nakayama, K., Rahman, M.T., Rahman, M., Katagiri, H., Nakayama, N., Ishikawa, M., Ishibashi, T., Iida, K., Kobayashi, H., et al. (2012). Loss of ARID1A expression is related to shorter progression-free survival and chemoresistance in ovarian clear cell carcinoma. Modern pathology : an official journal of the United States and Canadian Academy of Pathology, Inc 25, 282-288.

Kim, H., George, E., Ragland, R., Rafial, S., Zhang, R., Krepler, C., Morgan, M., Herlyn, M., Brown, E., and Simpkins, F. (2017). Targeting the ATR/CHK1 Axis with PARP Inhibition Results in Tumor Regression in BRCA-Mutant Ovarian Cancer Models. Clinical cancer research : an official journal of the American Association for Cancer Research 23, 3097-3108.

Kittler, R., Surendranath, V., Heninger, A.K., Slabicki, M., Theis, M., Putz, G., Franke, K., Caldarelli, A., Grabner, H., Kozak, K., et al. (2007). Genome-wide resources of endoribonuclease-prepared short interfering RNAs for specific loss-of-function studies. Nature methods 4, 337-344.

Knezevic, C.E., Wright, G., Remsing Rix, L.L., Kim, W., Kuenzi, B.M., Luo, Y., Watters, J.M., Koomen, J.M., Haura, E.B., Monteiro, A.N., et al. (2016). Proteome-wide Profiling of Clinical PARP Inhibitors Reveals Compound-Specific Secondary Targets. Cell chemical biology 23, 1490-1503.

Lehar, J., Krueger, A.S., Avery, W., Heilbut, A.M., Johansen, L.M., Price, E.R., Rickles, R.J., Short, G.F., 3rd, Staunton, J.E., Jin, X., et al. (2009). Synergistic drug combinations tend to improve therapeutically relevant selectivity. Nature biotechnology 27, 659-666.

Liu, X., Shi, Y., Maag, D.X., Palma, J.P., Patterson, M.J., Ellis, P.A., Surber, B.W., Ready, D.B., Soni, N.B., Ladror, U.S., et al. (2012). Iniparib nonselectively modifies cysteine-containing proteins in tumor cells and is not a bona fide PARP inhibitor. Clinical cancer research : an official journal of the American Association for Cancer Research 18, 510-523.

Lord, C.J., and Ashworth, A. (2013). Mechanisms of resistance to therapies targeting BRCA-mutant cancers. Nature medicine 19, 1381-1388.

Martins, M.M., Zhou, A.Y., Corella, A., Horiuchi, D., Yau, C., Rakshandehroo, T., Gordan, J.D., Levin, R.S., Johnson, J., Jascur, J., et al. (2015). Linking tumor mutations to drug responses via a quantitative chemical-genetic interaction map. Cancer discovery 5, 154-167.

McCabe, N., Turner, N.C., Lord, C.J., Kluzek, K., Bialkowska, A., Swift, S., Giavara, S., O’Connor, M.J., Tutt, A.N., Zdzienicka, M.Z., et al. (2006). Deficiency in the repair of DNA damage by homologous recombination and sensitivity to poly(ADP-ribose) polymerase inhibition. Cancer research 66, 8109-8115.

Mitchison, T.J. (2012). The proliferation rate paradox in antimitotic chemotherapy. Molecular biology of the cell 23, 1-6.

Mohni, K.N., Thompson, P.S., Luzwick, J.W., Glick, G.G., Pendleton, C.S., Lehmann, B.D., Pietenpol, J.A., and Cortez, D. (2015). A Synthetic Lethal Screen Identifies DNA Repair Pathways that Sensitize Cancer Cells to Combined ATR Inhibition and Cisplatin Treatments. PloS one 10, e0125482.

Muellner, M.K., Uras, I.Z., Gapp, B.V., Kerzendorfer, C., Smida, M., Lechtermann, H., Craig-Mueller, N., Colinge, J., Duernberger, G., and Nijman, S.M. (2011). A chemical-genetic screen reveals a mechanism of resistance to PI3K inhibitors in cancer. Nature chemical biology 7, 787-793.

Murai, J., Huang, S.Y., Das, B.B., Renaud, A., Zhang, Y., Doroshow, J.H., Ji, J., Takeda, S., and Pommier, Y. (2012). Trapping of PARP1 and PARP2 by Clinical PARP Inhibitors. Cancer research 72, 5588-5599.

Murai, J., Huang, S.Y., Renaud, A., Zhang, Y., Ji, J., Takeda, S., Morris, J., Teicher, B., Doroshow, J.H., and Pommier, Y. (2014). Stereospecific PARP trapping by BMN 673 and comparison with olaparib and rucaparib. Molecular cancer therapeutics 13, 433-443.

Norquist, B., Wurz, K.A., Pennil, C.C., Garcia, R., Gross, J., Sakai, W., Karlan, B.Y., Taniguchi, T., and Swisher, E.M. (2011). Secondary somatic mutations restoring BRCA1/2 predict chemotherapy resistance in hereditary ovarian carcinomas. Journal of clinical oncology : official journal of the American Society of Clinical Oncology 29, 3008-3015.

Olaussen, K.A., Dunant, A., Fouret, P., Brambilla, E., Andre, F., Haddad, V., Taranchon, E., Filipits, M., Pirker, R., Popper, H.H., et al. (2006). DNA repair by ERCC1 in non-small-cell lung cancer and cisplatin-based adjuvant chemotherapy. The New England journal of medicine 355, 983-991.

Shen, J., Peng, Y., Wei, L., Zhang, W., Yang, L., Lan, L., Kapoor, P., Ju, Z., Mo, Q., Shih Ie, M., et al. (2015). ARID1A Deficiency Impairs the DNA Damage Checkpoint and Sensitizes Cells to PARP Inhibitors. Cancer discovery 5, 752-767.

Shen, Y., Rehman, F.L., Feng, Y., Boshuizen, J., Bajrami, I., Elliott, R., Wang, B., Lord, C.J., Post, L.E., and Ashworth, A. (2013). BMN 673, a novel and highly potent PARP1/2 inhibitor for the treatment of human cancers with DNA repair deficiency. Clinical cancer research : an official journal of the American Association for Cancer Research 19, 5003-5015.

Subramanian, A., Tamayo, P., Mootha, V.K., Mukherjee, S., Ebert, B.L., Gillette, M.A., Paulovich, A., Pomeroy, S.L., Golub, T.R., Lander, E.S., et al. (2005). Gene set enrichment analysis: a knowledge-based approach for interpreting genome-wide expression profiles. Proceedings of the National Academy of Sciences of the United States of America 102, 15545-15550.

Swisher, E.M., Lin, K.K., Oza, A.M., Scott, C.L., Giordano, H., Sun, J., Konecny, G.E., Coleman, R.L., Tinker, A.V., O’Malley, D.M., et al. (2017). Rucaparib in relapsed, platinum-sensitive high-grade ovarian carcinoma (ARIEL2 Part 1): an international, multicentre, open-label, phase 2 trial. The Lancet Oncology 18, 75-87.

Yokoyama, Y., Matsushita, Y., Shigeto, T., Futagami, M., and Mizunuma, H. (2014). Decreased ARID1A expression is correlated with chemoresistance in epithelial ovarian cancer. Journal of gynecologic oncology 25, 58-63.

